# Doxorubicin inhibits phosphatidylserine decarboxylase and confers broad-spectrum antifungal activity

**DOI:** 10.1101/2023.04.07.535992

**Authors:** Yaru Zhou, Juan Zhao, Lei Yang, Ruiqing Bi, Ziting Qin, Peng Sun, Renjian Li, Mengfei Zhao, Yin Wang, Guang Chen, Hu Wan, Lu Zheng, Xiao-Lin Chen, Guanghui Wang, Qiang Li, Guotian Li

## Abstract

- As phospholipids of cell membranes, phosphatidylethanolamine (PE) and phosphatidylserine (PS) play crucial roles in glycerophospholipid metabolism. Broadly, some phospholipid biosynthesis enzymes serve as potential fungicide targets. Therefore, revealing the functions and mechanism of PE biosynthesis in plant pathogens would provide potential targets for crop disease control.
- We performed analyses including phenotypic characterizations, lipidomics, enzyme activity, site-directed mutagenesis, and chemical inhibition assays to study the function of PS decarboxylase-encoding gene *MoPSD2* in rice blast fungus *Magnaporthe oryzae*.
- The *Mopsd2* mutant was defective in development, lipid metabolism and plant infection. The PS level increased while PE decreased in *Mopsd2*, consistent with the enzyme activity. Furthermore, chemical doxorubicin inhibited the enzyme activity of MoPsd2 and showed antifungal activity against ten phytopathogenic fungi including *M. oryzae* and reduced disease severity of two crop diseases in the field. Three predicted doxorubicin-interacting residues are important for MoPsd2 functions.
- Our study demonstrates that MoPsd2 is involved in *de novo* PE biosynthesis and contributes to the development and plant infection of *M. oryzae* and that doxorubicin shows broad-spectrum antifungal activity as a fungicide candidate. The study also implicates that bacterium *Streptomyces peucetius*, which biosynthesizes doxorubicin, could be potentially used as an eco-friendly biocontrol agent.

## Introduction

Fungal diseases of crops, including rice blast and Fusarium head blight (FHB) of wheat, pose serious threats to global food security(Pennisi, 2010). Annual worldwide losses resulted from fungal diseases have been estimated to be 125 million dollars (Fisher, *et al*., 2012). Among different disease control strategies, application of fungicides is an important means to reduce the losses and improve the yield and quality of agricultural products (Russell, 2005).

Rice blast caused by the filamentous ascomycete, *Magnaporthe oryzae*, affects worldwide production of rice and causes yield losses up to 30% (Wilson & Talbot, 2009; Dean, *et al*., 2012). To initiate plant infection, the conidium of *M. oryzae* germinates and then develops a dome-shaped structure called the appressorium (Ebbole, 2007). During appressorium maturation, lipid droplets and glycogen migrate to the appressorium to provide nutrients and energy. For plant infection, the enormous turgor pressure that accumulates in mature appressoria is used to directly penetrate the plant cell wall (Howard, *et al*., 1991), and then the invasive hyphae grow and extend to neighboring cells as infection develops (Mosquera, *et al*., 2009). In the infection process, the fungus has to evade or suppress host reactive oxygen species (ROS) (Gao, *et al*., 2021). Finally, large numbers of conidia are produced from the disease lesions and released into the air to initiate a new disease cycle when conditions are favorable. Because *M. oryzae* has a very short infection cycle and sporulates abundantly, its pathogenesis and resistance to chemicals evolve dynamically. Accordingly, identification of novel targets for fungal disease control is required.

Phospholipids, essential components of biological membranes, play important roles in maintaining the cell structure and signal transduction (Dowhan, 1997; Rella, *et al*., 2016). Phosphatidylethanolamine (PE) and phosphatidylcholine (PC) are two main phospholipids, accounting for over half of the total phospholipids in eukaryotes(Vance & Steenbergen, 2005). The biosynthesis of PE and PC involves two pathways, the *de novo* pathway and Kennedy pathway (Fig. **S1**) (Carman & Han, 2011). In the *de novo* pathway, cytidine-diphosphate diacylglycerol is converted into phosphatidylserine (PS) that is subsequently decarboxylated by PS decarboxylases (Psds) into PE, and PE through three consecutive methylations by PEN-methyltransferases (PEMTs) to biosynthesize PC (Birner, *et al*., 2001). In the Kennedy pathway, ethanolamine and choline are converted into PE and subsequentially PC (Gibellini & Smith, 2010).

Enzymes involved in phospholipid biosynthesis play crucial roles in the development and pathogenesis of several pathogens (Wang, *et al*., 2019). The model unicellular eukaryote *Saccharomyces cerevisiae* contains two Psds, Psd1 and Psd2, and deletion of both Psds leads to impaired growth and defects in mitochondrial respiration (Trotter, *et al*., 1995; Birner, *et al*., 2001). In the human fungal pathogen *Candida albicans*, the *psd1Δ/Δ psd2Δ/Δ* mutant is severely defective in PE biosynthesis and growth (Chen, *et al*., 2010). Psd2 plays crucial roles in growth, morphogenesis and phospholipid homeostasis in the model filamentous fungus *Aspergillus nidulans* (Takagi, *et al*., 2021). In the wheat fungal pathogen *Fusarium graminearum*, the causal agent of Fusarium head blight (FHB), Psd2 is involved in sexual and asexual reproduction, lipid droplet formation, autophagy, and pathogenesis (Tang, *et al*., 2021). Some phospholipid biosynthesis proteins have been used as agrochemical targets (Pan, *et al*., 2018). Moreover, doxorubicin, typically produced by the bacterium *Streptomyces peucetius*, induces lipid peroxidation and suppresses Psd activities (Niraula, *et al*., 2010; Bellance, *et al*., 2020). However, involvement of Psds in pathogenicity and the possible inhibitory effects of doxorubicin on Psds in *M. oryzae* have not been studied.

Here, we showed that deletion of *MoPSD2* in *M. oryzae* resulted in defects in hyphal growth, conidiation, conidial morphology, lipid and glycogen utilization, and cell wall integrity. During plant infection, the *Mopsd2* mutant triggered over-accumulated ROS in rice and was attenuated in virulence. Moreover, doxorubicin substantially inhibited enzyme activities of MoPsd2 and showed antifungal activity to ten fungal pathogens. The three predicted doxorubicin-interacting residues are important for MoPsd2 functions. In summary, this study demonstrates that MoPsd2 plays vital roles in infection-related morphogenesis and virulence in *M. oryzae*, and reveals a broad-spectrum fungicide candidate, doxorubicin, that inhibits Psd2 enzyme activities, fungal development and plant infection.

## Materials and Methods

### Strains and culture conditions

The *M. oryzae* wild-type (WT) strains P131 and Guy11 and related transformants were routinely cultured on oatmeal agar (OMA) plates at 28°C (Chen, *et al*., 2014). The growth rate and conidiation were measured on minimal medium (MM) and OMA plates, respectively, at 5 days post-inoculation (dpi) (Ding, *et al*., 2010; Li, GT, *et al*., 2017). Transformants were selected on Terrific Broth3 (TB3) plates supplemented with 250 μg ml^-1^ of hygromycin B (Genview, USA) or 400 μg ml^-1^ of geneticin (GLPBIO, USA).

For stress tolerance assays, fungal strains were cultured at 28°C on MM plates supplemented with 1 M sorbitol or 1 mM ethanolamine, and on CM plates supplemented with 0.1 mg ml^-1^ Calcofluor white (CFW), 0.01% sodium dodecyl sulfate (SDS), or 5 mM H_2_O_2_ (Park, *et al*., 2006; Chen, *et al*., 2010).

Fungal strains stored in the laboratory were used to determine the concentration of doxorubicin that inhibits vegetative growth of these strains (Li, *et al*., 2023). Potato dextrose agar (PDA) plates supplemented with 0.15 mM doxorubicin were used to culture other fungal pathogens, including *Botrytis cinerea* B05.10, *Bipolaris maydis* TM17, *Bipolaris sorokiniana* LanKao9-3, *F. graminearum* PH-1, *Leptosphaeria biglobosa* E18-14, *Monilinia fructicola* 2YFT2-2, *Valsa mali* VM-1, and *Valsa pyri* VA-1 at 28℃. *Ustilaginoidea virens* strain JS60-2 was cultured on potato sucrose agar (PSA) plates. All strains used in this study are described in Table **S1**.

### Targeted gene deletion and complementation

The split-marker method was used to generate the *Mopsd2* mutant as previously described (Catlett, *et al*., 2003). Briefly, 1.1-kb upstream and 1.8-kb downstream flanking fragments of *MoPSD2* were amplified with primer pairs *PSD2*-1F/2R and *PSD2*-3F/4R, respectively. We used primers HYG-F/R to amplify the *hph* cassette from plasmid pCB1003. The flanking fragments that overlapped the *hph* cassette were transformed into the WT protoplasts (Yu, *et al*., 2004). Putative knockout mutants were verified by PCR.

To generate the *MoPSD2* complementation strain, the *MoPSD2* gene including its native 1.5-kb promoter was amplified with primers *MoPSD2*-CF/R and cloned into plasmid pKNTG (Chen, *et al*., 2014). *MoPSD2^N75A^*, *MoPSD2^D111A^* and *MoPSD2^D129A^*mutant alleles of *MoPSD2* were generated using the same yeast GAP repair approach as previously described (Bourett, *et al*., 2002). The construct was sequenced, and the correct one was transformed into protoplasts of *Mopsd2*. Primers used are listed in Table **S2**.

### Staining assays

Conidial suspensions (1×10^5^ conidia ml^-1^) were inoculated onto hydrophobic coverslips for 2, 4, 8, 12, and 24 h, and then stained with the Nile Red solution (Sigma-Aldrich, Germany) and KI/I_2_ solution, respectively, for lipid droplets and glycogen in germ tubes and appressoria (Thines, *et al*., 2000; Kimura, *et al*., 2004). The stained samples were examined under an epifluorescence microscope (Nikon Ni90, Japan).

Barley leaves inoculated with the WT and *Mopsd2* mutant strains at 36 hours post-inoculation (hpi) were stained with the Metal Enhanced DAB Substrate Kit (Solarbio, China) for 8 h and de-stained with ethanol/acetic acid (24:1, v/v) for 9 h. The infected leaves were then examined and photographed under the microscope (Nikon Ni90, Japan).

Plant infection assays with *M. oryzae*

Plant infection assays were performed as previously described (Valent, *et al*., 1991). Briefly, conidial suspensions of the WT (1×10^5^ conidia ml^-1^) were spray-inoculated on one-week-old rice seedlings of cultivar LTH (*Oryza sativa* cv. Lijiangxintuanheigu) and drop-inoculated on five-day-old barley seedlings of cultivar E9 (*Hordeum vulgare* cv. E9) (Yang, *et al*., 2022). Symptoms on rice and barley were photographed at 7 and 5 dpi, respectively. In addition, conidial suspensions of the WT (1×10^5^ conidia ml^-1^) were punch-inoculated on four-week-old rice seedlings of LTH as previously described (Li, WT, *et al*., 2017). Briefly, 10-μl spore suspension was inoculated on wounded rice leaves, and the infected plants were kept in the greenhouse. Symptoms were examined at 14 dpi.

To assay the penetration rate, conidial suspensions (1.2×10^5^ conidia ml^-1^) were drop-inoculated (8 μl) onto the surface of five-day-old barley epidermis and incubated in a dark chamber at 28℃. The infection level was examined at 30 hpi under a microscope (Nikon Ni90, Japan).

### Protein purification, western blotting and enzyme activity analyses

Expression of MoPsd2-6×His was performed as previously described (You, *et al*., 2022). The complementary DNA (cDNA) of *MoPSD2* was cloned into vector pET-28a, and the construct was introduced into *E. coli* strain BL21 (DE3). The resultant *E. coli* strain was cultured in Luria-Bertani (LB) to reach the optical density (OD_600_) at 0.6-0.8 and then induced with 1 mM Isopropyl β-D-Thiogalactoside (IPTG) for 12 h. The total proteins were isolated and the His-tagged protein was purified with Ni NTA Beads 6FF (Smart-Lifesciences Biotechnology, China) as previously described (Raran-Kurussi & Waugh, 2017).

For western blotting assays, mycelia were harvested from two-day-old liquid CM cultures at 28℃ and total proteins were extracted as previously described (Li, *et al*., 2011). We used 10% SDS-PAGE gel to separate proteins. The Western Blotting Detection Kit (Advansta, USA) was used to visualize the target signal with ChemiDoc Touch Imaging System (BIO-RAD, USA). For the protein loading control, the gel was stained with coomassie brilliant blue (CBB) and washed with the de-staining buffer before examination (Wang, *et al*., 2018).

The Psd enzyme activity assay based on fluorescence was performed with 1,2-diacetyl benzene/β-mercaptoethanol (1,2-DAB/β-ME) (Sigma-Aldrich, Germany), which forms a fluorescent iso-indole-mercaptide conjugate with PE, as previously described (Choi, *et al*., 2020). Briefly, enzyme activity assays were performed with a series of concentrations of purified MoPsd2-6×His, 0.5 mM PS, and 0.2% Triton X-100 in a buffer of NaCl (50 mM) and potassium phosphate (10 mM, pH 7.4) at 30℃ for 60 min. Potassium phosphate (10 mM, pH 9.85) was added to the solution to terminate the reaction. Then 61 mM 1,2-DAB and 5% β-ME was added and the sample was incubated at 22℃ for 60 min. The standard curve was made with PE (0 to 0.5 mM). The data were performed in four independent experiments. PE (0.25 mM) with or without doxorubicin (0.04 mM) were used as controls. Fluorescence intensity was monitored at the excitation spectrum (λ_ex_) = 364 nm and the emission spectrum (λ_em_) = 425 nm with the SPARK 10M multimode reader (TECAN, Austria).

### Subcellular localization assays

The MoPsd2-GFP construct was transformed into the WT protoplasts as previously described (Chen, *et al*., 2014). Transformants were screened with the primer pair *PSD2*-5F/pKNT-seqR, and verified for fluorescence signals under the microscope (Nikon Ni90, Japan). The confirmed transformants were observed for GFP fluorescence signals in conidia, appressoria and hyphae under a confocal microscope Leica TCS SP8 (Leica Microsystems, Germany). The λ_ex_ for the GFP was set at 498 nm and the λ_em_ at 571 nm. The images were collected and analyzed with a 40×objective lens using LAS X software.

### Lipidomics analysis

Mycelia of the WT and *Mopsd2* mutant were harvested from two-day-old liquid CM cultures at 28℃. Lipid extraction was followed the method as described (Bligh & Dyer, 1959; Hanson & Lester, 1980). Briefly, mycelial samples were freeze-dried and incubated in hot isopropanol (75°C) containing 0.05% butylated hydroxytoluene (BHT) (Sigma-Aldrich, Germany) for 15 min. Chloroform/methanol (2:1, v/v) with 0.01% BHT was used to extract the total lipids. Then 1 M KCl and water were sequentially added to the sample. The upper phase was discarded after centrifugation. All supernatants were pooled and dried under a nitrogen gas flow. According to the dry weight of mycelia, chloroform was proportionally added to the sample in 10 mg ml^-1^. Phospholipids were measured in an Agilent HPLC system together with the 4000 QTRAP MS system (Applied Biosystems, USA) as previously described (Shen, *et al*., 2010).

To separate different phospholipids, two-dimensional thin layer chromatography (2D-TLC) was performed on TLC plates (Macherey Nagel, Germany) (Zhao, *et al*., 2022). First, the TLC plate was preheated at 110°C for 90 min. Chloroform/methanol/ammonium hydroxide (65: 25: 5, v/v/v) and chloroform/methanol/acetic-acid/water (85: 15: 10: 3, v/v/v/v) were used for the first- and second-dimensional separation, respectively. The plates were dried for 20 min and then exposed to iodine vapor for 90 seconds to visualize the lipid spots.

### Fungal growth, sexual reproduction and infection assays with doxorubicin

Conidial germination and appressorium formation of *M. oryzae* with or without doxorubicin were assayed on hydrophobic glass coverslips (Liu, *et al*., 2011). Briefly, conidial suspensions were drop-inoculated onto hydrophobic glass coverslips, and then incubated in a moist chamber at 28℃. The percentage of germination was examined under the microscope at 2 h and the appressorium formation at 24 h. For sexual reproduction assays, *F. graminearum* was cultured on carrot agar plates supplemented with or without 0.15 mM doxorubicin. Starting at 5 dpi, 800 μl Tween 20 (0.1%) was added to each plate, and aerial hyphae were pressed down every day. Homothallic perithecium formation was examined at 15 dpi (Bowden & Leslie, 1999). PDA plates with or without 0.15 mM doxorubicin were used to culture *B. cinerea* and *S. sclerotiorum* at 20℃ for sclerotium formation assays (Li, *et al*., 2018). The number of mature sclerotia from each plate was examined at 10 dpi.

To analyze the possible inhibitory effect of doxorubicin on different pathogens, conidia of various species, including *M. oryzae* P131, *B. sorokiniana* LanKao9-3, *B. maydis* TM17, *B. cinerea* B05.10, and *V. pyri* VA-1 with or without 0.15 mM doxorubicin were inoculated on leaves of barley, wheat, maize, and pear and fruits of tomato with methods described in previous studies, respectively (Ding, *et al*., 2015; Sarven, *et al*., 2020; Nie, *et al*., 2021; Zhang, WY, *et al*., 2022). Symptoms were examined and analyzed at 5 dpi.

### Protein structure prediction and molecular docking

The protein sequence of MoPsd2 was from NCBI (https://www.ncbi.nlm.nih.gov/) and the protein structure of MoPsd2 was predicted with Alphafold2 (Bryant, *et al*., 2022). The chemical structure of doxorubicin was from the PubChem Compound database (https://www.ncbi.nlm.nih.gov/pccompound/). Molecular docking of MoPsd2 with doxorubicin was performed with Autoduck (Forli, *et al*., 2016).

### RNA-seq analysis

Mycelia of the WT with or without 0.1 mM doxorubicin treatment and the *Mopsd2* strains were cultured in liquid CM with shaking at 150 rpm, 28℃ for 48 h (Tang, *et al*., 2020). Total RNA was extracted with a FastPure Plant Total RNA Isolation Kit (Vazyme, China) and quality-controlled. The sequencing was performed on the BGISEQ-500 sequencer at Nextomics Biosciences (Wuhan, China). Clean reads were aligned to the *M. oryzae* MG8 genome (http://fungi.ensembl.org/index.html) with HISAT2_2.2.1 after filtering the adapter and low-quality reads (Huang, *et al*., 2017). FeatureCounts_2.0.1 was used to count the read numbers, and DESeq2 for gene expression analysis (Love, *et al*., 2014). Genes with |log2 fold change| ≥ 1 and false discovery rate ≤ 0.05 were defined as differentially expressed genes (DEGs), which were further analyzed for the KEGG pathway enrichment (https://www.keggjp/) (Yu, *et al*., 2012).

### RT-qPCR assays

Mycelia used for RT-qPCR assays were the same as the samples used in the RNA-seq analysis. The 1^st^ Strand cDNA Synthesis Kit (Vazyme, China) was used to generate cDNA. RT-qPCR assays were performed on the CFX Connect real-time PCR system (BioRad, America) with TransStart Tip Green qPCR SuperMix (TransGen Biotech, China). The *M. oryzae* actin gene (MGG_00604) was used as the internal control. The relative gene expression level was calculated by the 2^-ΔΔCt^ method (Livak & Schmittgen, 2001). Primers used are listed in Table **S2**.

### Phytotoxicity assays

Possible phytotoxicity of doxorubicin was assessed on four-week-old rice seedlings of cultivar CO39 and two-week-old wheat seedlings of cultivar Huamai 8 (Wipfler, *et al*., 2019). Doxorubicin was sprayed onto rice at 0.5 mM and onto wheat at 0.3-mM. The fresh weight of rice and wheat seedlings was examined at 7 and 5 dpi, respectively. For lipidomics assays, doxorubicin (0.15 mM, 0.5 mM) was applied to 14-day-old rice plants. Seven days later, the treated rice leaves were harvested for analyses.

### Field trials

For field trials, one-month-old seedlings of rice cultivar CO39 were transplanted to the blast nursery in Yangjiang, Guangdong Province in March as previously described (Liang, *et al*., 2021). The field trials used a randomized plot method with three replicates, each with 600 seedlings in 30 m^2^. Thirty days after planting, 0.5 mM doxorubicin was sprayed onto the seedlings and repeated once three days later. Water was used as the control. The disease index was examined at 16 dpi.

To determine the effect of doxorubicin on FHB, field experiments were performed on flowering wheat heads of cultivar Xiaoyan22 at Yangling, Shaanxi Province (Han, *et al*., 2020). Conidia of the *F. graminearum* strain PH-1 were collected (1×10^5^ conidia ml^-1^) from cultures in the carboxymethyl cellulose medium (CMC) supplemented with 0.3 mM doxorubicin for inoculation assays at 10 μl per wheat head. Conidial suspensions with sterile water served as the control. Twenty biological replicates were used for each treatment. The disease index was examined and plants were photographed at 14 dpi.

## Results

### Characterization of MoPsd1 and MoPsd2 in *M. oryzae*

To determine whether MoPsd1 and MoPsd2 proteins are conserved among different species, we identified MoPsd1 (MGG_09213) and MoPsd2 (MGG_07037) in the genome of *M. oryzae* and compared them with homologs from different organisms, including *A. nidulans*, *B. cinerea*, *C. albicans*, *F. graminearum*, *Fusarium oxysporium*, *Fusarium verticillioides*, *Neurospora crassa*, *S. cerevisiae*, *Schizosaccharomyces pombe*, and *Sclerotinia sclerotiorum*. Phylogenetic analysis showed that both MoPsd1 and MoPsd2 are highly conserved, and among the homologs analyzed, NcPsd1 and NcPsd2 show the highest homology with MoPsd1 and MoPsd2, respectively (Fig. **S2**). *MoPSD2* encodes an 805-residue protein, with a PS-decarboxylase and two C2 domains.

### The *Mopsd2* mutant is reduced in growth and conidiation, and hypersensitive to stressors

To elucidate the function of *MoPSD1* and *MoPSD2* in *M. oryzae*, we attempted to delete the two genes by homologous recombination (Fig. **S3a**). After screening more than 200 hygromycin-resistant transformants from four independent experiments, we failed to obtain the null mutant for *MoPSD1*. For *MoPSD2*, two *Mopsd2* knockout transformants were confirmed by PCR (Fig. **S3b**). For genetic complementation, the *MoPSD2* gene, including its native promoter was amplified, cloned and transformed into protoplasts of the *Mopsd2* mutant.

To analyze the role of *MoPSD2* in fungal development, we cultured *Mopsd2* on different media (Fig. **1a**). Growth rates of the *Mopsd2* mutant on OMA, CM and MM plates were decreased by 8%, 10% and 53%, respectively, compared to the WT (Fig. **1b**). The average cell length at the hyphal tips of the *Mopsd2* mutant was 37% shorter than that of the WT (Fig. **S4a, b**), indicating that *MoPSD2* plays an important role in vegetative growth. Conidiation assays showed that deletion of *MoPSD2* reduced conidia production by approximately 84% (Fig. **1c**). In addition, 70% of the *Mopsd2* conidia had three cells, and the other 30% one or two cells, in contrast to 93% three-celled conidia in the WT (Fig. **S4c**). These results indicated that the loss of *MoPSD2* affects fungal growth, conidiation and conidial morphology.

**Fig. 1.**
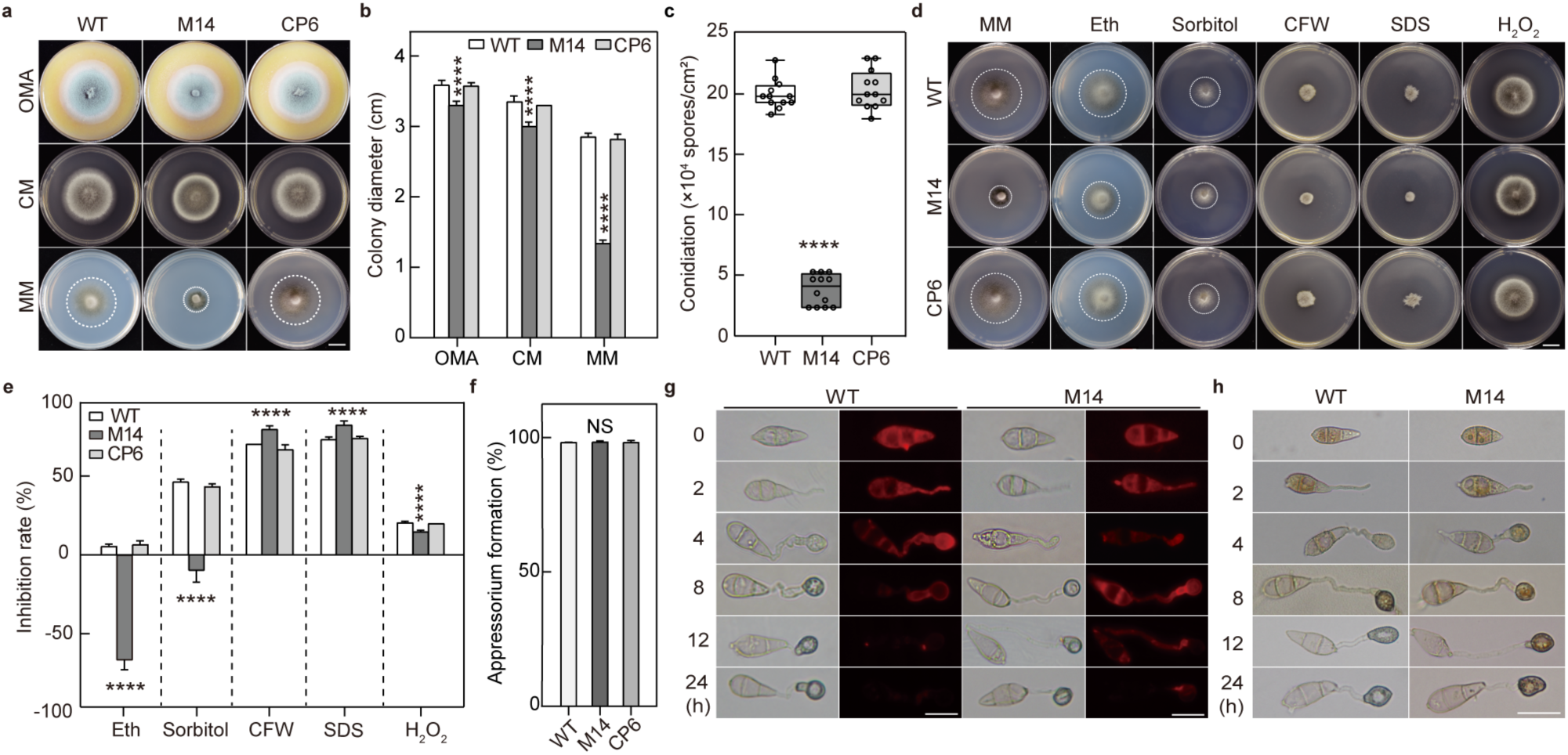
MoPsd2 is involved in vegetative growth, conidiation, lipid and glycogen utilization, and stress responses. (**a**) Five-day-old oatmeal medium (OMA), complete medium (CM) and minimal medium (MM) cultures of the wild-type (WT), *Mopsd2* mutant (M14) and *Mopsd2*/MoPsd2-GFP complementation (CP6) strains at 28℃. Bar, 1 cm. (**b**) Colony diameters of the strains shown in (**a**) were measured and calculated from seven biological replicates. (**c**) Conidiation of the WT, M14 and CP6 strains from five-day-old OMA cultures. (**d**) Colony morphology of the WT, M14 and CP6 strains supplemented with different stressors on MM or CM plates. CFW, Calcofluor white; Eth, ethanolamine; SDS, sodium dodecyl sulfate. Bar, 1 cm. (**e**) Inhibition by different stressors on the same set of strains shown in (**d**) at 5 days post-inoculation (dpi). Means and standard deviations were calculated from three biological replicates. (**f**) Appressorium formation of the WT, M14 and CP6 strains on hydrophobic coverslips. (**g**) Lipid droplets were stained with Nile Red during appressorium formation. Bars, 25 μm. (**h**) Glycogen of the WT and M14 strains was stained with the iodine solution. Bar, 25 μm. Data were analyzed with Student’s *t*-test. Asterisks represent significant differences (\*\*\*\**P* < 0.0001); NS, not significant.

To determine whether *MoPSD2* is involved in stress responses in *M. oryzae*, the WT, *Mopsd2* mutant and complementation strains were cultured on CM and MM plates supplemented with different stressors (Fig. **1d**). Growth inhibition rates of the *Mopsd2* mutant were increased by 13% and 12%, respectively, under cell wall-disturbing reagents 0.1 mg ml^-1^ of CFW and 0.01% SDS compared to the WT. Furthermore, the growth rates of the *Mopsd2* mutant under the osmotic stressor sorbitol (1 M) and ethanolamine (1 mM) were increased, while the growth rates of the WT and complementation strains decreased under the same conditions (Fig. **1e**). Thus, *MoPSD2* plays differential roles in responses to different stressors.

### *MoPSD2* is involved in utilization of lipid and glycogen and in plant infection

Appressoria are important for *M. oryzae* infection. We analyzed appressorium formation of the mutant and observed that the *Mopsd2* mutant was normal in appressorium formation (Fig. 1**f**). To analyze the utilization of nutrients during appressorium formation, lipid droplets and glycogen were stained with Nile red and I_2_/KI solutions, respectively. In the WT strain, the lipid droplets began to degrade at 8 hpi, and almost no corresponding fluorescence was detected at 24 hpi. In contrast, fluorescence signals indicating lipid droplets were still observed in germ tubes and appressoria at 24 hpi in the *Mopsd2* mutant (Fig. **1g**). Glycogen was transferred from the conidium to the appressorium in the WT strain at 8 hpi, but in the *Mopsd2* mutant strain, the glycogen was enriched in the conidia at 12 hpi (Fig. **1h**). These results indicated that *MoPSD2* plays an important role in degradation and utilization of lipid droplets and glycogen in *M. oryzae*.

In plant infection assays, compared to lesion areas developed on leaves inoculated with the WT and complementation strains (Fig. **2a, b, c**), the lesion areas on leaves inoculated with the *Mopsd2* mutant in punch-, drop- and spray-inoculation assays decreased by 51%, 43% and 66%, respectively (Fig. **2d, e**), indicating that *MoPSD2* plays an indispensable role in full virulence of *M. oryzae*. We found that the infection rate in barley epidermal cells by the *Mopsd2* mutant was reduced by 20% (Fig. **2f**). To investigate the possible reason of attenuated virulence of the *Mopsd2* mutant and analyze whether *MoPSD2* is involved in circumventing host ROS, barley leaves were infected with the WT and *Mopsd2* mutant strains and stained with 3,3’-diaminobenzidine (DAB) at 30 hpi (Fig. **2g**). When observed under the microscope, nearly 80% of cells infected by the *Mopsd2* mutant formed brown precipitates, indicating ROS accumulation, but only 29% of the cells infected with the WT strain showed brown precipitates (Fig. **2h**). The results indicated that *MoPSD2* is involved in either evading host immunity or suppressing host ROS for full fungal virulence.

**Fig. 2.**
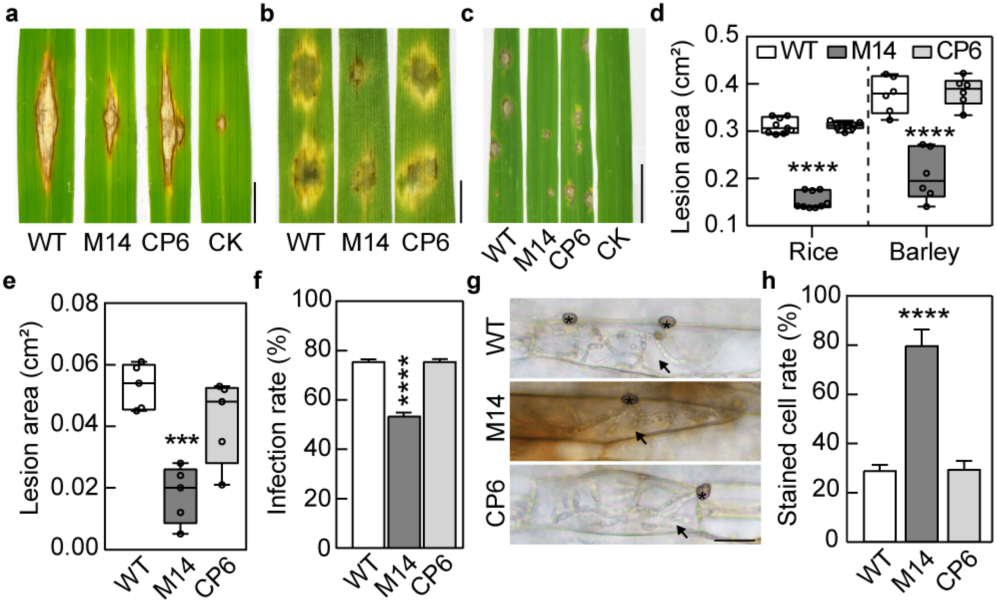
Deletion of *MoPSD2* leads to attenuated virulence and increased ROS accumulation in plant cells. (**a**) Punch-inoculation of rice leaves with the wild-type (WT), *Mopsd2* mutant (M14) and *Mopsd2*/MoPsd2-GFP complementation (CP6) strains, and CK (0.025% Tween 20). Symptoms were examined at 14 dpi. Bar, 0.5 cm. (**b**) Drop-inoculation assays on detached barley leaves with conidial suspensions (1×10^5^ conidia ml^-1^) at 5 dpi. Bar, 0.5 cm. (**c**) Seedlings of rice cultivar LTH were spray-inoculated with WT, M14, CP6, and CK. Photos were taken at 7 dpi. Bar, 0.5 cm. (**d**) Lesion areas on rice and barley leaves by the same set of strains shown in (**b**). (**e**) Lesion areas on rice by the same set of strains shown in (**c**). (**f**) Infection rates of WT, M14 and CP6 strains in barley epidermal cells. (**g**) DAB staining of the infected barley cells at 36 hours post-inoculation (hpi). Asterisks and arrows indicate appressoria and infectious hyphae, respectively. Bar, 20 μm. (**h**) Percentages of stained barley cells were determined at 8 hpi. Data were analyzed with Student’s *t*-test. Asterisks represent significant differences (\*\*\**P* < 0.001, \*\*\*\**P* < 0.0001).

### MoPsd2 catalyzes phosphatidylserine (PS) into phosphatidylethanolamine (PE)

Before analyzing the biochemical function of MoPsd2, we analyzed the subcellular localization of MoPsd2 during the development of *M. oryzae.* The *MoPSD2*-GFP construct was transformed into the WT. The GFP signals appeared mostly in the cytoplasm in conidia, appressoria and hyphae (Fig. **3a**).

**Fig. 3.**
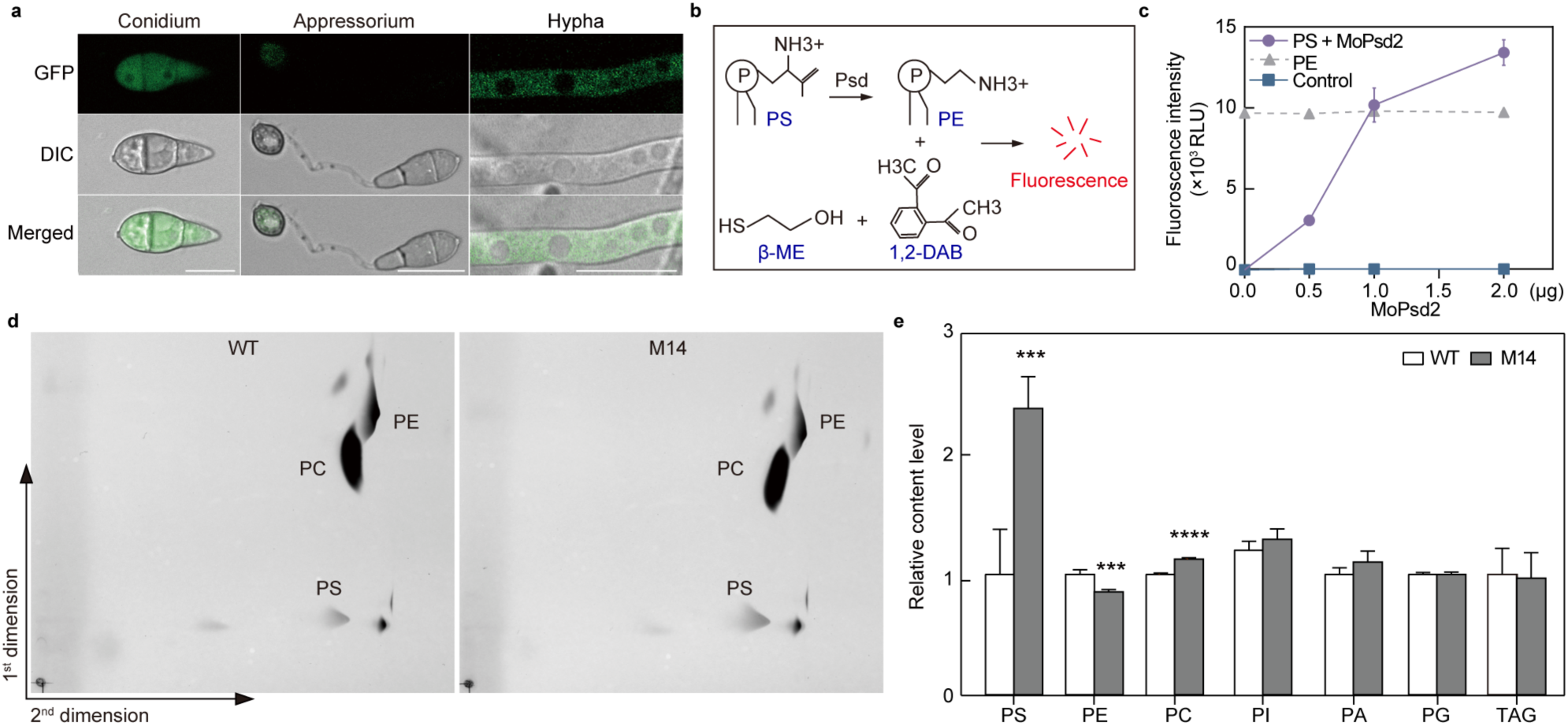
MoPsd2 is involved in the conversion of phosphatidylserine (PS) into phosphatidylethanolamine (PE) in *M. oryzae*. (**a**) Subcellular localization of MoPsd2-GFP at different developmental stages. Bars, 20 μm. (**b**) Schematic diagram of fluorescence-based assays for Psd activity, that is, PE detection with 1,2-diacetyl benzene/β-mercaptoethanol (1,2-DAB/β-ME). (**c**) Fluorescence-based enzyme activity assays of MoPsd2. Heat-inactivated MoPsd2 and PE were used as the control. RLU, relative light unit. (**d**) Two-dimensional thin layer chromatography (2D-TLC) of phospholipids isolated from the wild-type (WT) and *Mopsd2* (M14) strains. PC, phosphatidylcholine. (**e**) Lipidomics assays were performed on five biological replicates. PA, phosphatidic acid; PG, phosphatidylglycerol; PI, phosphatidylinositol; TAG, triacylglycerol. Data were analyzed with Student’s *t*-test. Asterisks represent significant differences (\*\*\**P* < 0.001, \*\*\*\**P* < 0.0001).

For enzyme activity assays, MoPsd2-6×His was expressed in *Escherichia coli* and purified (Fig. **S5**). We used a fluorescence-based Psd assay to analyze the enzyme activities by detecting fluorescence from the PE conjugate (Fig. **3b**) (Choi, *et al*., 2020). The fluorescence signals were detected only when MoPsd2 was added to the reaction. The results indicated that MoPsd2 converts PS into PE (Fig. **3c**).

To further examine the function of MoPsd2 in *vivo*, we performed 2D-TLC and lipidomics assays. The 2D-TLC analysis showed that the PS content in the *Mopsd2* mutant was higher than the WT control (Fig. **3d**). The lipidomics analysis further confirmed that the levels of PS and PC were increased by 138% and 13%, respectively, while PE was reduced by 15% in the *Mopsd2* mutant compared to that in the WT (Fig. **3e**). Taken together, these results show that MoPsd2 is involved in *de novo* biosynthesis of PE in *M. oryzae*.

### Doxorubicin suppresses growth, development and plant infection of *M. oryzae*

We investigated the effect of doxorubicin, a putative Psd inhibitor, on fungal development in *M. oryzae*. On MM plates supplemented with 0.08 mM doxorubicin, the growth rate of *M. oryzae* was slightly reduced by 15% (Fig. **4a**), while conidiation was seriously reduced by 72% (Fig. **4b**). The concentration for 50% of maximal effect (EC50) of doxorubicin was 1.152 mM based on the inhibition rate of mycelial growth on MM plates (Fig. **S8**). Moreover, when conidial suspensions were incubated with 0.08 mM doxorubicin, only 60% conidia germinated and 90% of those failed to form appressoria (Fig. **4c**). In plant infection assays, the lesion area and lesion number decreased as the concentration of doxorubicin increased (Fig. **4d, e**), and lesions barely developed on barley leaves inoculated with conidial suspensions in the presence of doxorubicin at concentration of 0.06 mM or higher (Fig. **4e**). Invasive hyphae formed in epidermal cells of barley leaves were restricted by doxorubicin (Fig. **4f**). These results demonstrated that doxorubicin suppresses fungal development and plant infection of *M. oryzae*.

**Fig. 4.**
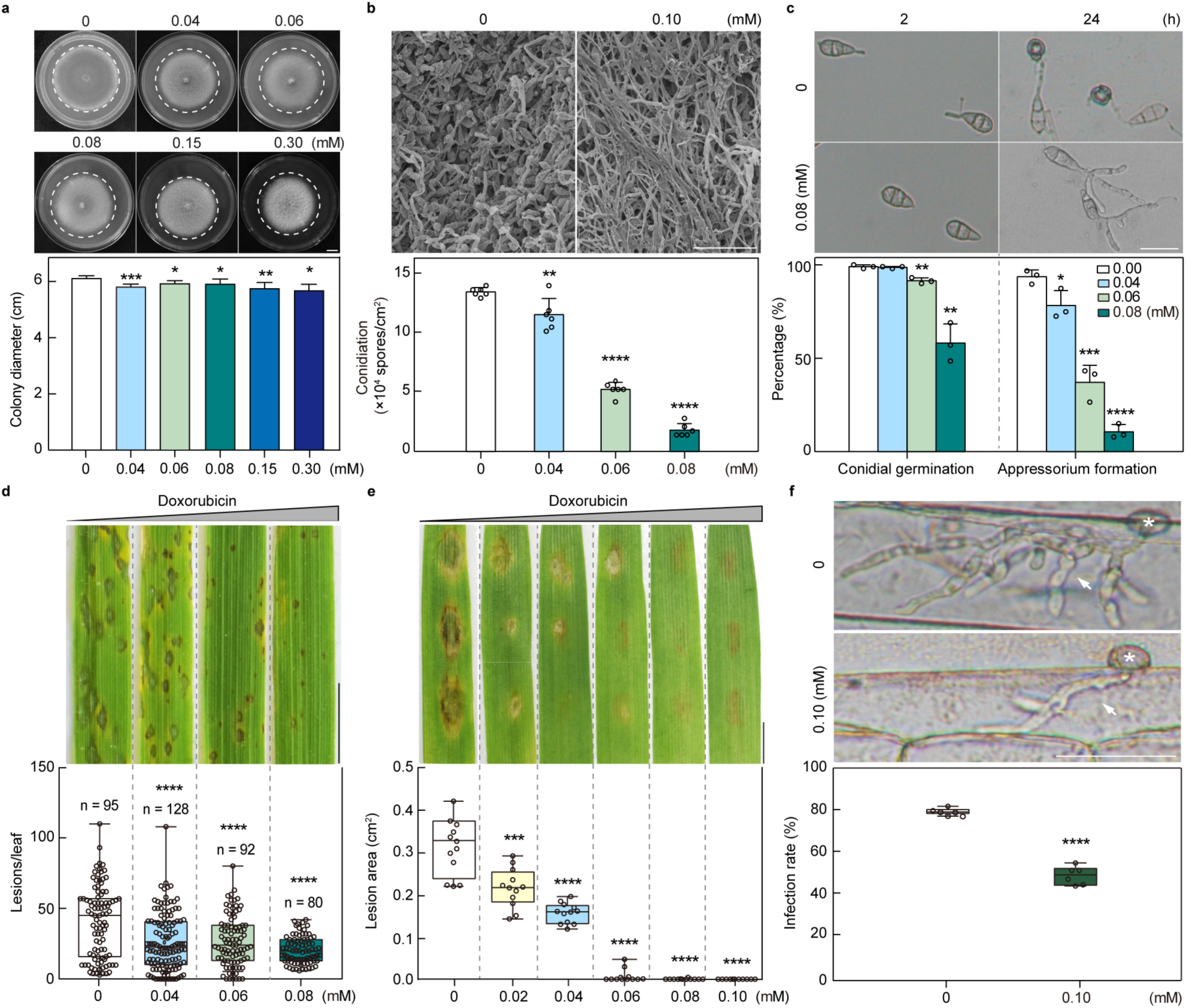
Doxorubicin inhibits fungal growth, development and plant infection by *M. oryzae*. (**a**) Twelve-day-old MM cultures supplemented with 0, 0.04-, 0.06-, 0.08-, 0.15-, and 0.30-mM doxorubicin. Bar, 1 cm. Inhibitory effects of doxorubicin on mycelial growth of the wild-type (WT) were calculated from four biological replicates. (**b**) Scanning electron microscope (SEM) and conidiation assays under doxorubicin treatment. Bar, 50 μm. Five-day-old CM cultures with six biological replicates were used for conidiation assays. (**c**) Conidial germination (2 h) and appressorium formation (24 h) assays of the WT with or without doxorubicin. Conidial suspensions of the WT with different concentrations of doxorubicin were inoculated on hydrophobic glass coverslips. Bar, 20 μm. (**d**) Spray-inoculation assays on rice cultivar CO39 with conidial suspensions of *M. oryzae* strain Guy11 (1×10^5^ conidia ml^-1^) supplemented with different concentrations of doxorubicin. Bar, 0.5 cm. The lesion number was examined at 7 dpi. (**e**) Drop-inoculation assays on detached barley leaves with conidial suspensions of the WT (1×10^5^ conidia ml^-1^) in different concentrations of doxorubicin. Bar, 0.5 cm. The lesion area was examined at 5 dpi. (**f**) Penetration assays with or without doxorubicin. Asterisks and arrows indicate appressoria and invasive hyphae, respectively. Invasive hyphae developed in barley epidermal cells were examined at 30 hpi. Bar, 30 μm. The formation of invasive hyphae in barley epidermal cells were suppressed by doxorubicin. Data were analyzed with Student’s *t*-test. Asterisks indicate significant differences (\*\**P* < 0.01, \*\*\**P* < 0.001, \*\*\*\**P* < 0.0001).

### Doxorubicin inhibits MoPsd2 activities and alters lipid biosynthesis in *M. oryzae*

To investigate the connection between the MoPsd2 and doxorubicin, the molecular docking using the MoPsd2 structure predicted by Alphafold2 was performed and the results showed that amino acid residues Asn75, Asp111 and Asp129 of MoPsd2 potentially interact with doxorubicin (Fig. **5a**). To investigate whether doxorubicin inhibits the enzyme activity of MoPsd2, doxorubicin was added to the reaction buffer. The result showed that the enzyme activity of MoPsd2 was reduced by 86.3% in the presence of 0.04 mM doxorubicin (Fig. **5b**). In addition, we used lipidomics approaches to analyze the role of doxorubicin in the lipid biosynthesis pathway in *vivo*. The PS level in the doxorubicin-treated sample was two-fold higher than the control, consistent with the results in the *Mopsd2* mutant (Fig. **5c**).

**Fig. 5.**
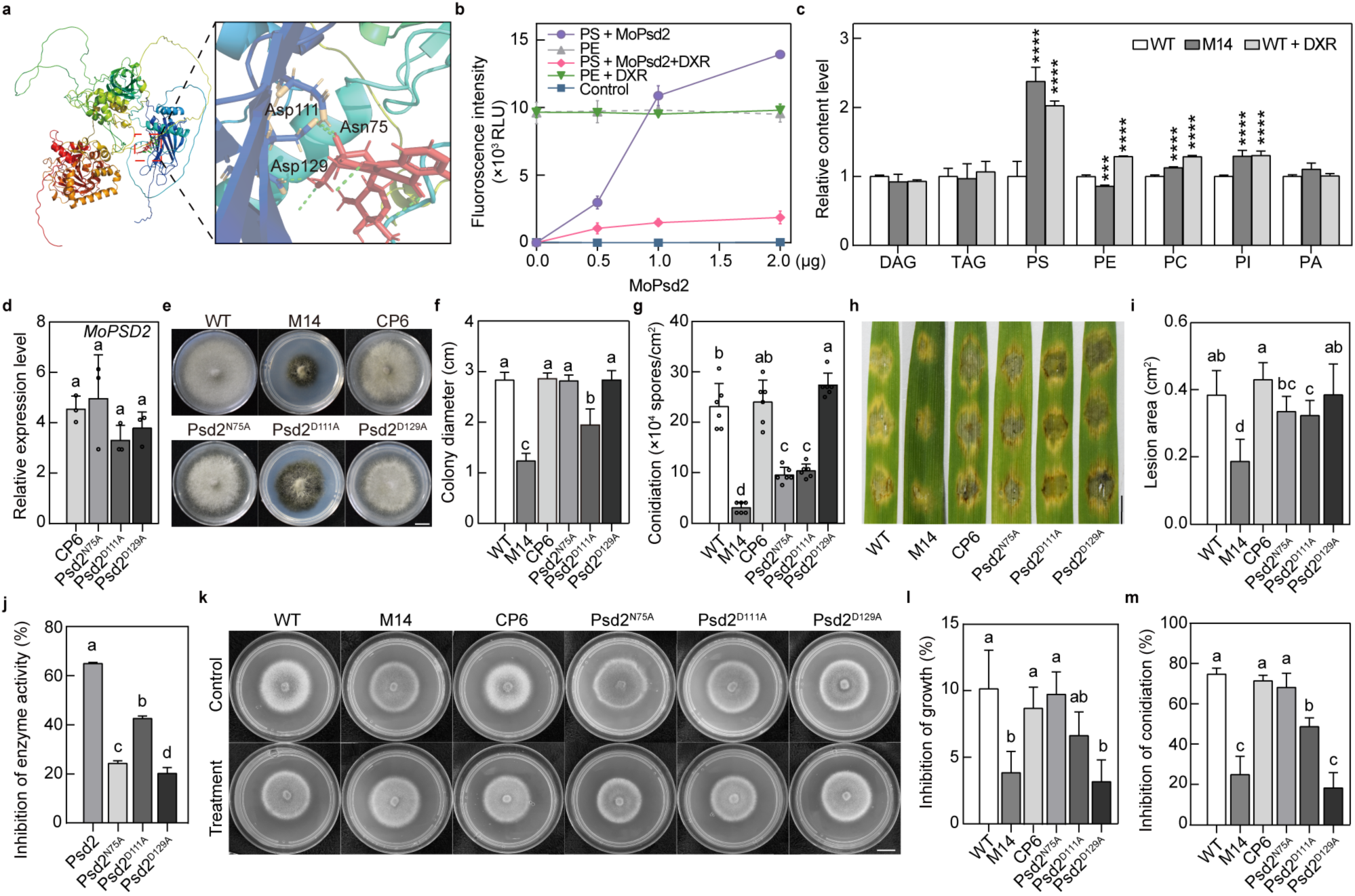
Doxorubicin inhibits MoPsd2 activity and alters lipid biosynthesis in *M. oryzae*. (**a**) Protein structure of MoPsd2 was predicted with AlphaFold2. Molecular docking of MoPsd2 with doxorubicin was simulated with Autodock. Amino acids Asn75, Asp111 and Asp129 of MoPsd2 in the inset represent predicted residues binding to doxorubicin (DXR). (**b**) The enzyme activities of MoPsd2 were suppressed by doxorubicin. The heat-inactivated MoPsd2 protein, PE with or without doxorubicin were used as controls. RLU, relative light unit. (**c**) Lipidomics assays of the wild-type (WT) strain cultured in the liquid CM supplemented with or without doxorubicin and the *Mopsd2* mutant (M14). Means and standard deviations were calculated from five biological replicates. Data were analyzed with Student’s t-test. Asterisks represent significant differences (\*\*\**P* < 0.001, \*\*\*\**P* < 0.0001). (**d**) Relative gene expression levels of *MoPSD2* in CP6, Psd2^N75A^, Psd2^D111A^, and Psd2^D129A^ strains. A, alanine; D, aspartate; N, asparagine. (**e**) Five-day-old MM cultures of the different strains. Colony diameters (**f**) and conidiation **(g)** of the strains from five-day-old OMA cultures. **(h)** Drop-inoculation assays on detached barley leaves with conidial suspensions (1×10^5^ conidia ml^-1^) at 5 dpi. Bar, 0.5 cm. (**i**) Lesion areas on barley leaves shown in (**h**). **(j)** Inhibition rates of the enzyme activity of MoPsd2, MoPsd2^N75A^, MoPsd2^D111A^, and MoPsd2^D129A^ by doxorubicin. **(k)** Five-day-old CM cultures supplemented with/without doxorubicin. Inhibition rates of vegetative growth (**l**) and conidiation (**m**) by doxorubicin calculated from five-day-old CM cultures. Data were analyzed using a one-way ANOVA with Duncan’s new multiple range test.

To analyze the function of predicted doxorubicin-binding amino acids of MoPsd2, the predicted residues were substituted and the resultant mutant alleles were transformed into the *Mopsd2* mutant. The expression levels of different *MoPSD2* mutant alleles in the resultant strains were comparable to that in the complementation strain, higher than that in the WT (Fig. 5**d**). We investigated vegetative growth, conidiation and virulence, and the results show that *MoPSD2^N75A^* fully rescued the growth defect of the *Mopsd2* mutant but not conidiation or virulence. Furthermore, the *MoPSD2^D111A^* allele did not completely rescue defects in fungal growth, conidiation or virulence of the mutant. These results demonstrate that these residues play important and varied roles in the function of MoPsd2 (Fig. **5e-i**).

Complementarily, the *in vitro* enzymatic activity assays showed that doxorubicin effectively suppresses the activity of MoPsd2, but the inhibition rates for the site-directed mutants were reduced (Fig. **5j**). Similar results were observed in hyphal growth and conidiation of the strains expressing these mutant alleles with doxorubicin (Fig. **5k, l, m**). In these alleles, *MoPSD2^D129A^* completely rescued the biological phenotypes of the *Mopsd2* mutant, but was significantly reduced for the inhibition under doxorubicin treatment, comparable to the level of the null mutant. We speculate that the Asp129 residue may be the predicted binding site of the doxorubicin to MoPsd2, though not that important for the function of MoPsd2.

RNA-seq and reverse transcription quantitative PCR (RT-qPCR) assays were performed to investigate the possible mechanism and signaling pathways that doxorubicin was involved in. Out of 366 DEGs, 298 were up-regulated and 68 down-regulated with doxorubicin treatment (Fig. **S6a**). Non-overlapping gene clusters were derived from the *Mopsd2* mutant and *M. oryzae* treated with or without doxorubicin, consisting of up-regulated genes and down-regulated genes both in the *Mopsd2* mutant and doxorubicin-treated groups (Fig. **S6d, e**). KEGG enrichment analysis showed that the mitogen-activated protein kinase (MAPK) pathway and lipid metabolism were negatively regulated both in the doxorubicin-treated and *Mopsd2* mutant groups (Fig. **S6f**). RT-qPCR assays showed that expression levels of *MoPSD2* was reduced to 84% in the doxorubicin-treated group (Fig. **S9**) (Carman & Han, 2011). DEGs resulted from doxorubicin treatment were categorized into four groups, representing genes involved in growth, conidiation, appressorium formation, and pathogenesis (Fig. **S6b, c**) (Badaruddin, *et al*., 2013; Lu, *et al*., 2014; Huang, *et al*., 2022). Taken together, those results indicated that doxorubicin inhibits the enzyme activities of MoPsd2, leading to increased PS levels, and alters expression levels of genes involved in development, lipid biosynthesis and pathogenesis in *M. oryzae*.

### Doxorubicin shows broad-spectrum antifungal activity

Because Psd2 is a target of doxorubicin in *M. oryzae*, we analyzed Psd2 homologs in ten plant pathogens and found that Psd2 was conserved in different fungi (Fig. **6a**). In addition, predicted protein structures of these Psd2 homologs were close to the structure of MoPsd2, and the closest BsPsd2 had a template modeling score (TM-score) of 0.64 (Fig. **S10**). To investigate the effect of doxorubicin on vegetative growth of these pathogens, PDA plates supplemented with 0.15 mM doxorubicin were used to culture different fungal pathogens, including *B. cinerea*, *B. maydis*, *B. sorokiniana, F. graminearum*, *L. biglobosa*, *M. fructicola, U. virens, V. mali*, and *Valsa pyri*. *M. oryzae* was cultured on MM plates supplemented with 0.15 mM doxorubicin (Fig. **6b**). The growth inhibition rates of those ten pathogens by doxorubicin ranged from 7% to 45%, with *V. pyri* affected the most. In addition, we selected *B. cinerea* and *F. graminearum* for conidial germination assays and found that conidial germination of the two fungi was almost completely suppressed by 0.15 mM doxorubicin (Fig. **6c**). In addition, sexual reproduction was inhibited as the number of perithecia produced from the doxorubicin-treated *F. graminearum* culture was reduced by 34% (Fig. **6d**). The sclerotium formation of *B. cinerea* was almost completely suppressed and that for *S. sclerotiorum* was decreased by 46% by 0.15 mM doxorubicin (Fig. **6e**). These results indicate that doxorubicin effectively inhibits growth and development of multiple fungal pathogens.

**Fig. 6.**
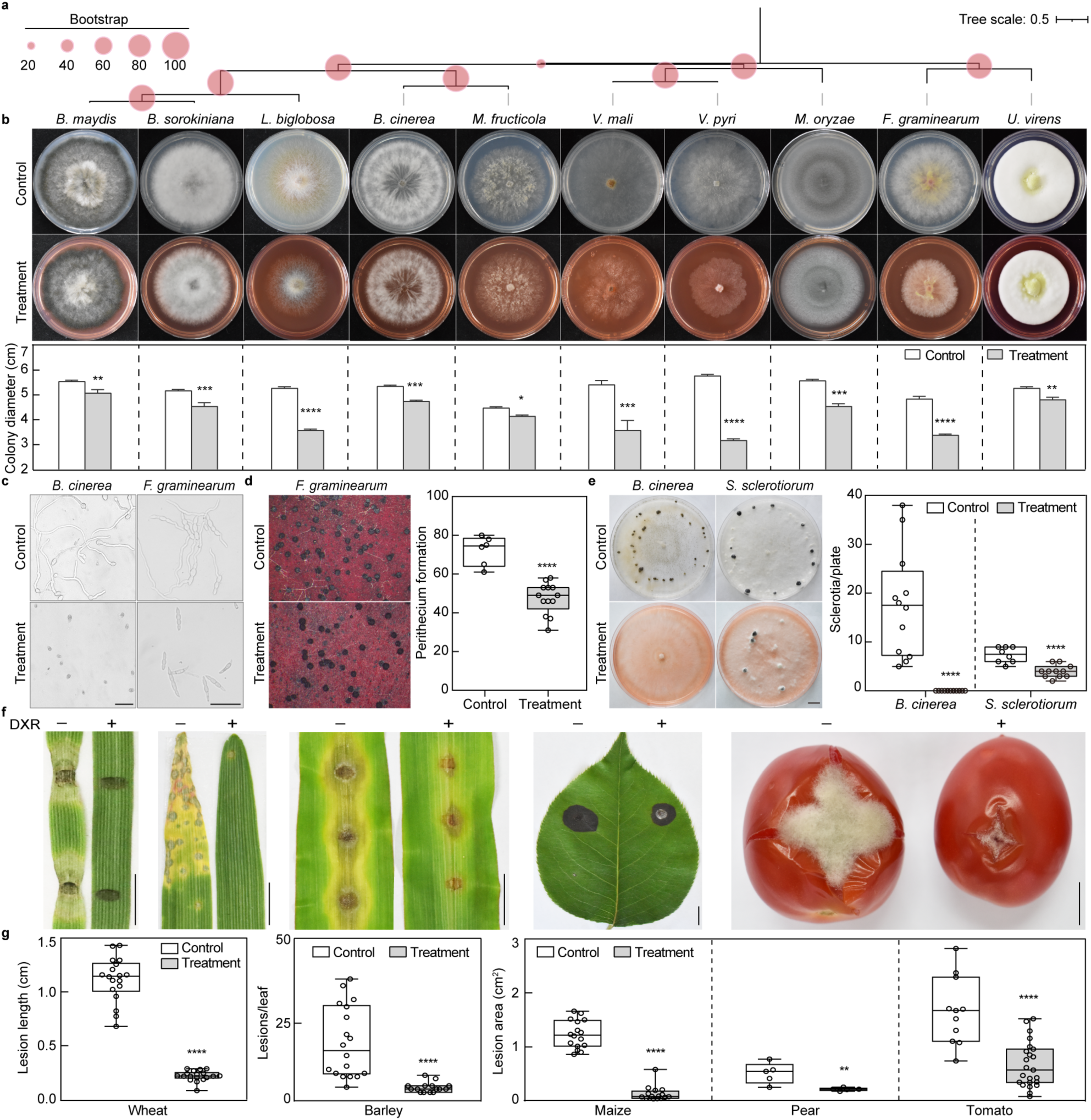
Doxorubicin inhibits development and plant infection of multiple fungal pathogens. (**a**) Phylogenetic tree of Psd2 homologs from ten fungal phytopathogens. (**b**) Doxorubicin inhibited hyphal growth of ten phytopathogenic fungi. Colony morphology of strains grown on plates supplemented with 0.15 mM doxorubicin. (**c**) Doxorubicin inhibited conidial germination of *B. cinerea* and *F. graminearum*. Bars, 50 μm. (**d**) Sexual production of *F. graminearum* was suppressed by doxorubicin. Fifteen-day-old carrot agar cultures were examined for the perithecium formation. (**e**) Doxorubicin inhibited sclerotium formation of *B. cinerea* and *S. sclerotiorum*. Black dots in plates indicate sclerotia. Bar, 1 cm. The number of sclerotia was examined at 10 dpi. (**f**) Plant infection assays of fungal strains supplemented with or without 0.15 mM doxorubicin (DXR). Bars, 1 cm. (**g**) The lesion length, number and area in different infection assays were examined at 5 dpi. Data were analyzed with Student’s *t*-test. Asterisks represent significant differences (\**P* < 0.05, \*\**P* < 0.01, \*\*\**P* < 0.001, \*\*\*\**P* < 0.0001).

Plant infection assays of *M. oryzae* on barley, *B. maydis* on maize, *V. pyri* on pear, *B. cinerea* on tomato, and *B. sorokiniana* on wheat (Fig. **6f**) showed that doxorubicin reduced disease symptoms by 63%-88%, as indicated by lesion length, area or number (Fig. **6g**). These results together showed that doxorubicin effectively inhibits plant infection by multiple plant pathogens.

### Doxorubicin is effective in the control of crop diseases in the field

The efficacy of doxorubicin in disease control was evaluated on rice blast and FHB of wheat in the field. Rice seedlings were planted in a blast nursery and treated with 0.5 mM doxorubicin (Fig. **7a**). The results showed that symptoms of rice blast in the doxorubicin treatment group were less serious than that in the control group. Compared to the water control, the disease index of rice blast of the doxorubicin-treated group was reduced by 20% (Fig. **7b**). Moreover, conidial suspensions of *F. graminearum* supplemented with or without 0.3 mM doxorubicin were used to inoculate wheat heads in the field (Fig. **7c**). The disease index of FHB was reduced by 44% with 0.3 mM doxorubicin treatment compared to the control (Fig. **7d**). These results demonstrate that doxorubicin efficiently reduces disease severity of rice blast and FHB in the field.

**Fig. 7.**
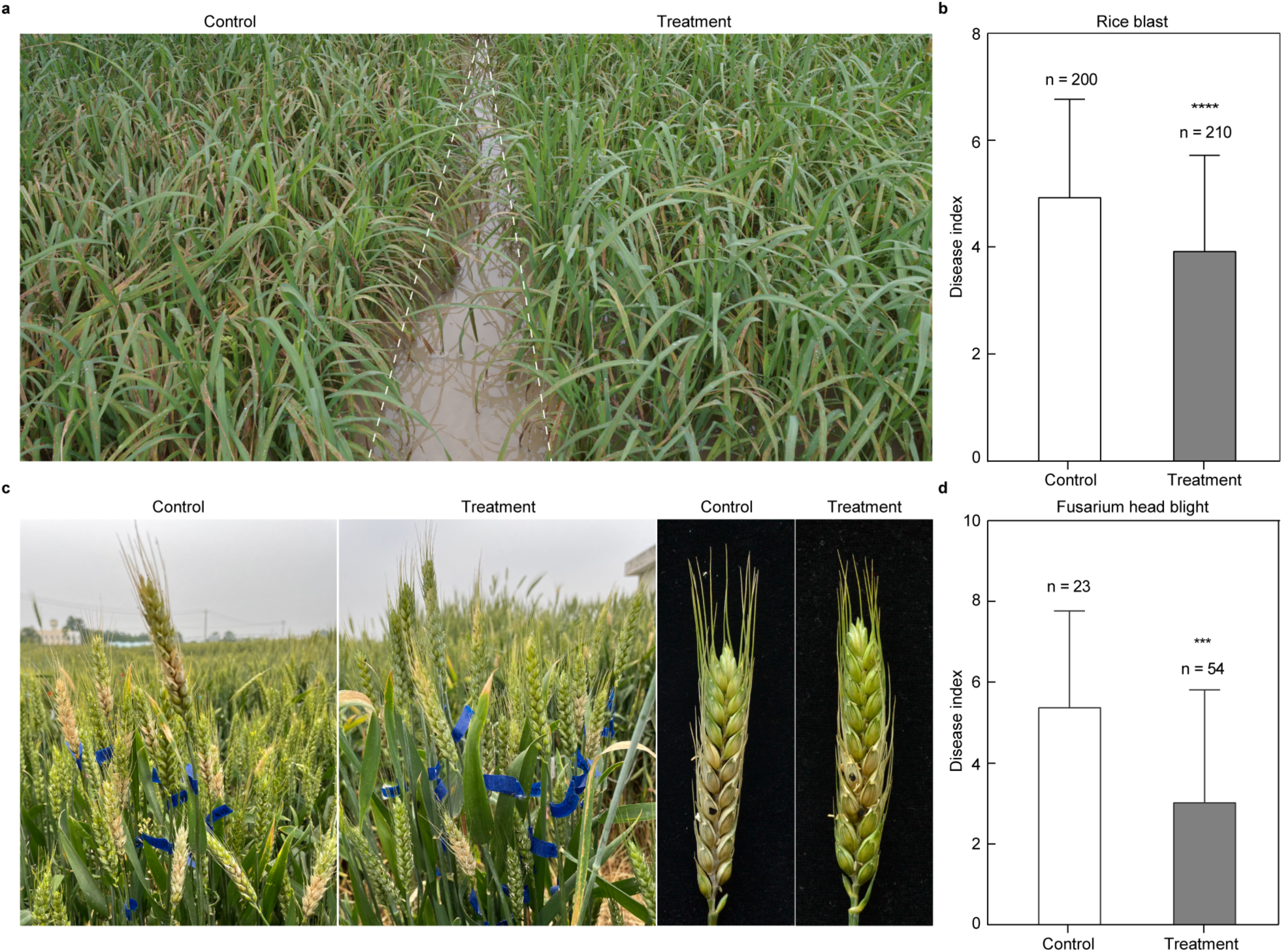
Doxorubicin reduces the severity of crop diseases in the field. (**a**) Field trials of doxorubicin on rice seedlings CO39 in the field. Doxorubicin (0.5 mM) or water was sprayed onto the rice seedlings after 30 days planting. (**b**) Disease index of rice blast were examined at 16 dpi. (**c**) Symptoms of Fusarium head blight of wheat. Conidial suspensions of *F. graminearum* strain PH-1 with or without doxorubicin (0.3 mM) were inoculated on wheat heads. (**d**) The disease index of Fusarium head blight was examined at 14 dpi. Means and standard deviations were calculated from the number of independent replicates as indicated. Data were analyzed with Student’s *t*-test. Asterisks represent significant differences (\*\*\**P* < 0.001, \*\*\*\**P* < 0.0001).

## Discussion

Psds, which convert PS into PE, are essential for maintaining cellular PE levels. In this study, we sought to functionally analyze MoPsd2 in the development and pathogenesis of *M. oryzae* and determine whether MoPsd2 may be used as a potential target for novel fungicides. Our results showed that MoPsd2 is involved in the regulation of vegetative growth, conidiation and virulence in *M. oryzae*. Furthermore, we demonstrated that doxorubicin, a natural metabolite of the filamentous soil bacterium *S. peucetius*, inhibited MoPsd2 activity and resulted in reduced fungal growth, infection-related morphogenesis and plant infection. Doxorubicin also showed broad-spectrum antifungal activity against ten fungal pathogens. These findings reveal functions of MoPsd2 in *M. oryzae* and provide a fungicide candidate for disease control.

The *Mopsd2* mutant was defective in hyphal growth and conidiation in *M. oryzae*, similar to the *psd2* mutant of *A. nidulans* (Takagi, *et al*., 2021), supporting the conserved role of *PSD2* in fungal development. In addition, mutants defective in PE biosynthesis show defects in cell wall integrity. The *Mopsd2* mutant and *psd1Δ/Δ psd2Δ/Δ* mutant of *C. albicans* are hypersensitive to membrane-perturbing agents, such as SDS (Wolf, *et al*., 2015). MoPsd2 expresses Psd activity (Fig. **3c**), and deletion of *MoPSD2* resulted in increased levels of PS and decreased levels of PE, similar to the changes observed in the yeast *psd2* mutant (Birner, *et al*., 2001). Interestingly, ethanolamine, a substrate for PE biosynthesis in an alternative pathway, partially rescued growth defects of the *Mopsd2* mutant, which was also observed in the *Fgpsd2* mutant (Wang, *et al*., 2019). Taken together, our results showed that Psds are involved in PE biosynthesis, fungal development and cell wall integrity. In contrast to the observations in the aforementioned fungi, deletion of *PSD2* in *S. cerevisiae* caused little or no defect in growth and stress responses, although some changes in phospholipid levels were demonstrated (Birner, *et al*., 2001). We observed that MoPsd2 is localized in the cytoplasm (Fig. **3a**), unlike the Golgi-localized yeast Psd2, which may be one reason for functional differences between MoPsd2 and yeast Psd2 (Trotter, *et al*., 1995). In summary, Psds are generally conserved in PE biosynthesis but subtly different roles may be operating in different fungi.

*MoPSD2* and other phospholipid biosynthesis-related genes play important roles in plant infection. The *Mopsd2* mutant either triggered abundant ROS production or failed to suppress host immunity and was significantly attenuated in virulence (Fig. **2**), consistent with the *psd2* mutant of *F. graminearum* (Tang, *et al*., 2021). Similarly, deletion of another phospholipid biosynthesis gene, *MoPAH1*, abolishes the formation of appressorium-like structures at the hyphal tip and consequently pathogenicity, and deletion of phospholipase C gene, *FgPLC*, disrupted conidial germination and virulence in *F. graminearum* (Ding, *et al*., 2015; Zhao, *et al*., 2022). Thus, the biosynthesis of phospholipids including PE plays an indispensable role in fungal pathogenesis. In *M. oryzae*, the involvement of PE biosynthesis in virulence may be through the regulation of the degradation and utilization of lipid droplets and glycogen during appressorium formation, which we showed are impaired in the *Mopsd2* mutant (Fig. **1g, h**) (Gok, *et al*., 2022). Deletion of *MoFAS1*, a gene encoding a fatty acid-degrading enzyme, results in defective melanin biosynthesis and impaired utilization of lipid droplets in *M. oryzae*, consistent with *Mopsd2* (Sangappillai & Nadarajah, 2020). In summary, deletion of *MoPSD2* causes perturbations of the phospholipid composition, which in turn delays lipid and glycogen metabolism among others, and attenuates fungal virulence.

Doxorubicin inhibited the enzyme activities of MoPsd2 (Fig. **5b**), indicating that MoPsd2 is one target of doxorubicin, consistent with results of a previous report showing that doxorubicin inhibits Psd activities and alters the PS/PE ratio in mammalian HeLa cells (Bellance, *et al*., 2020).

Doxorubicin-treated *M. oryzae* strains exhibit phenotypes reminiscent of the *Mopsd2* mutant, including reduced growth and conidiation, attenuated virulence and altered phospholipid levels. In accord, *GPH1*, *HOX7* and *CNF2*, which regulate conidiation, appressorium formation and plant infection in *M. oryzae* (Badaruddin, *et al*., 2013; Lu, *et al*., 2014; Huang, *et al*., 2022), are down-regulated by doxorubicin treatment. Additionally, the overlapped down-regulated genes from the *MoPsd2* mutant and doxorubicin-treated groups are enriched in the MAPK pathway and lipid metabolism, similar to that of *S. cerevisiae* (Taymaz-Nikerel, 2021). Residues Asn75, Asp111 and Asp129 of MoPsd2, which were predicted to interact with doxorubicin, are in the functionally conserved C2 domain (Fig. **S2**) (Kitamura, *et al*., 2002; Wu & Voelker, 2004). In consistence, when the first two residues were mutated, the resultant mutant alleles could not fully rescue the defects of the *Mopsd2* mutant. Additionally, the inhibition rate by doxorubicin on the *Mopsd2* mutant and Psd2^D129A^ strains were reduced, supporting that MoPsd2 is one target of doxorubicin in *M. oryzae* (Fig. **5**) and that Asp129 of MoPsd2 could be important for the interaction of doxorubicin with MoPsd2 (Qiu, *et al*., 2011; Vela-Corcía, *et al*., 2018). To summarize, doxorubicin inhibits development and plant infection of *M. oryzae* possibly through targeting the key residues in the conserved C2 domain of MoPsd2. In addition to MoPsd2, doxorubicin may have other targets of action, as the doxorubicin treatment on *M. oryzae* resulted in more phenotypes than that of the *Mopsd2* mutant. However, detailed inhibitory mechanisms of doxorubicin on *M. oryzae* need further investigation.

Doxorubicin effectively inhibited vegetative growth, conidial germination, sexual production, and plant infection in ten plant pathogens (Fig. **6**), consistent with the conserved Psd2 homologs in these pathogens. Broadly, phospholipids have diverse and critical roles in cellular functions, and relevant enzymes serve as potential targets for disease control. For example, the widely used fungicide, isoprothiolane, suppresses transmethylation in PC biosynthesis (Hu, *et al*., 2014).

Doxorubicin, targeting PE biosynthesis, showed inhibitory effects on rice blast and FHB in the field with no visible side effect on plant growth (Fig. **7** and Fig. **S7**). Our results show the potential application of doxorubicin as a fungicide for control of multiple plant fungal diseases, and other phospholipid biosynthesis related enzymes may be potential targets for fungicide development (Denning & Bromley, 2015).

In conclusion, MoPsd2, one enzyme involved in biosynthesis of PE, is involved in the development and infection-related morphogenesis in the devastating fungal pathogen *M. oryzae*. Additionally, doxorubicin inhibits the enzyme activities of MoPsd2, fungal growth and plant infection of *M. oryzae*. And the three predicted doxorubicin-interacting residues are important for MoPsd2 functions. Although these preliminary findings on broad-spectrum antifungal activities of doxorubicin are promising, additional research is needed to confirm and further solidify the results. Because the concentration for 50% of maximal effect of doxorubicin is relatively high (Fig. **S8**), chemical modification of doxorubicin is a promising route to improve its fungicidal efficiency (Madera-Santana, *et al*., 2018). Structure-based computational simulation, as a cutting-edge technology for drug discovery by virtually screening chemicals for the target protein, is a rigorous strategy to direct designs of doxorubicin derivatives to enhance its fungicidal efficiency (Fink, *et al*., 2022; Kaplan, *et al*., 2022). Alternatively, given that doxorubicin is a natural metabolite of bacterium *S. peucetius* and that the doxorubicin biosynthesis level can be significantly promoted in this industrial bacterium, it is worthwhile to investigate whether *S. peucetius* can be used as an eco-friendly biocontrol strain to control rice blast and a wide range of other devastating plant diseases (Noh, *et al*., 2010).

## Acknowledgements

We thank Drs. Larry D. Dunkle and Charles P. Woloshuk at Purdue University for critical reading of this manuscript and Ms. Dongqin Li from the National Key Laboratory of Crop Genetic Improvement for technical assistance in lipidomics analyses. The field trial on rice blast was assisted by Xianya Jiang at Yangjiang Institute of Agricultural Sciences. Maize seeds and pear leaves were kindly provided by Drs. Zhibin Lai and Guoping Wang, respectively, at Huazhong Agricultural University. We thank Huailong Teng and Huanyu Cai from college of Science of Huazhong Agricultural University for technical assistance. We thank all the lab members for helpful suggestions and discussions. This work was supported by National Key Research and Development Program of China (2022YFA1304402), Natural Science Foundation of China (32172373, 31801723) and Fundamental Research Funds for the Central Universities (2662020ZKPY006) to G. L. This work was also supported by Hubei Hongshan Laboratory.

## Author contributions

G.L. conceived and oversaw the project. Y.Z. initiated the designs and conducted biological experiments. L.Y. predicted the molecular docking, and R.B. and Z.Q. analyzed RNA-seq data. J.Z., P.S. and M.Z. participated in doxorubicin inhibition assays. R.L. collected some strains used in this study. Q.L. and G.C. performed the lipidomics assays. G.W. performed the field experiments on wheat. Y.W. performed the western blot assays. H.W., L.Z., and X.C. guided the paper writing and fruitful discussions. Y.Z. and G.L. wrote and edited the manuscript with inputs from all authors.

## Data availability

RNA-seq data used in this study are available at the National Genomics Data Center Genome Sequence Archive (https://ngdc.cncb.ac.cn/gsa/) under the accession number CRA008554.

