## supplemental Files for "Doxorubicin inhibits phosphatidylserine decarboxylase and confers broad-spectrum antifungal activity"

**Supporting Information**

**Table S1** Strains used in this study.

| Strain | Genotype | References |
|---|---|---|
| B05.10 | The wild-type strain of *Botrytis cinerea* | (Sarven, *et al.*, 2020) |
| TM17 | The wild-type strain of *Bipolaris maydis* | (Ding, *et al.*, 2015) |
| LanKao9-3 | The wild-type strain of *Bipolaris sorokiniana* | (Zhang, *et al.*, 2022) |
| PH-1 | The wild-type strain of *Fusarium graminearum* | (Ali, *et al.*, 2022) |
| E18-14 | The wild-type strain of *Leptosphaeria biglobosa* | (Cai, *et al.*, 2014) |
| 2YTF2-2 | The wild-type strain of *Monilinia fructicola* | (Chen, *et al.*, 2017) |
| P131 | The wild-type strain of *Magnaporthe oryzae* | (Peng & Shishiyama, 2011) |
| Psd2$^{D111A}$ | *MoPSD2*$^{D111A}$ in *Mopsd2* | This study |
| Psd2$^{D129A}$ | *MoPSD2*$^{D129A}$ in *Mopsd2* | This study |
| Psd2$^{N75A}$ | *MoPSD2*$^{N75A}$ in *Mopsd2* | This study |
| Guy11 | The wild-type strain of *M. oryzae* | (Zhang, *et al.*, 2018) |
| M14 | The *Mopsd2* deletion mutant of P131 | This study |
| M27 | The *Mopsd2* deletion mutant of P131 | This study |
| CP6 | Complementation transformant of *Mopsd2* | This study |
| AG-1 | The wild-type strain of *Rhizoctonia solani* | (Abdoulaye, *et al.*, 2017) |
| 1980 | The wild-type strain of *Sclerotinia sclerotiorum* | This study |
| JS60-2 | The wild-type strain of *Ustilaginoidea virens* | (Zhang, *et al.*, 2020) |
| VM-1 | The wild-type strain of *Valsa mali* | (Zhang, *et al.*, 2019) |
| VA-1 | The wild-type strain of *Valsa pyri* | This study |

**Table S2** Primers used in this study.

| Name | Sequence (5'-3') |
|---|---|
| *ACTIN*-qF | ccatgtaccctggtctttcg |
| *ACTIN*-qR | ttcgagatccacatctgctg |
| *CHO2*-qF | tcttcaacaaccgcaccaa |
| *CHO2*-qR | tgaaggcaagcgtgagtagg |
| *CPT1*-qF | gtcttcatcctgacgggctac |
| *CPT1*-qR | atcacattgcggcacgact |
| *EPT1*-qF | gccagccgttcccatact |
| *EPT1*-qR | gtcagggcgtttatctccaa |
| HYG-F | gtcgatgcgacgcaatcgt |
| HYG-R | gctgatctgaccagttgc |
| pKNT-seqR | gtgctgcttcatgtggtcgg |
| *PSD1*-qF | gcttccactcgccaactaa |
| *PSD1*-qR | cggtggtgagcgagtttgt |
| *PSD2*-CF | agggaacaaaagctgggtaccgcatcctgaagcatctgggt |
| *PSD2*-CR | gcccttgctcaccataagcttgcccgcaattctggttgtta |
| Psd2-His-F | ccggaattcatggttcgaataattcccagc |
| Psd2-His-R | cgcggatccgcccgcaattctggttgt |
| Psd2-m1F | atagtaccttgttttggcctgc |
| Psd2-m1R | gcaggccaaaacaaggtactat |
| Psd2-m2F | ctgctccttgccgtctgc |
| Psd2-m2R | gcagacggcaaggagcag |
| Psd2-m3F | cgagttcgcccttgcatta |
| Psd2-m3R | taatgcaagggcgaactcg |
| *PSD2*-qF | cttcttccaacggttccctc |
| *PSD2*-qR | tgttgatgccgtatgaccag |
| *PSD2*-1F | acttcttccccacgattccc |
| *PSD2*-2R | ttgacctccactagctccagccaagccgtgatgacagtggcgtttgg |
| *PSD2*-3F | gaatagagtagatgccgaccgcgggttctttgaggagggcaaga |
| *PSD2*-4R | ctgtcgtgatgattggaggg |
| *PSD2*-5F | ctcctcagcaggcgacagac |

| | |
|---|---|
| *PSD2*-6R | tgtgagggaaccgttggaag |
| *PSD2*-7F | caatgtcatcgtctcaagcc |
| *PSD2*-8R | gctgatctgaccagttgc |
| *PSD2*-9F | gtcgatgcgacgcaatcgt |
| *PSD2*-10R | aacacaatacgcaatgaggg |

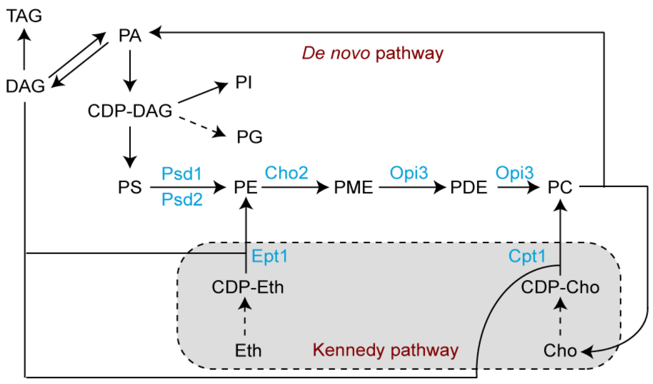


**Fig. S1** Schematic representation of major routes for phosphatidylethanolamine (PE) and phosphatidylcholine (PC) biosynthesis.

CDP-DAG, cytidine-diphosphate diacylglycerol; Cho, choline; Eth, ethanolamine; PA, phosphatidic acid; PDE, phosphatidyldimethylethanolamine; PG, phosphatidylglycerol; PI, phosphatidylinositol; PME, phosphatidylmonomethylethanolamine; PS, phosphatidylserine; TAG, triacylglycerol. The biosynthesis routes were adapted from yeast.

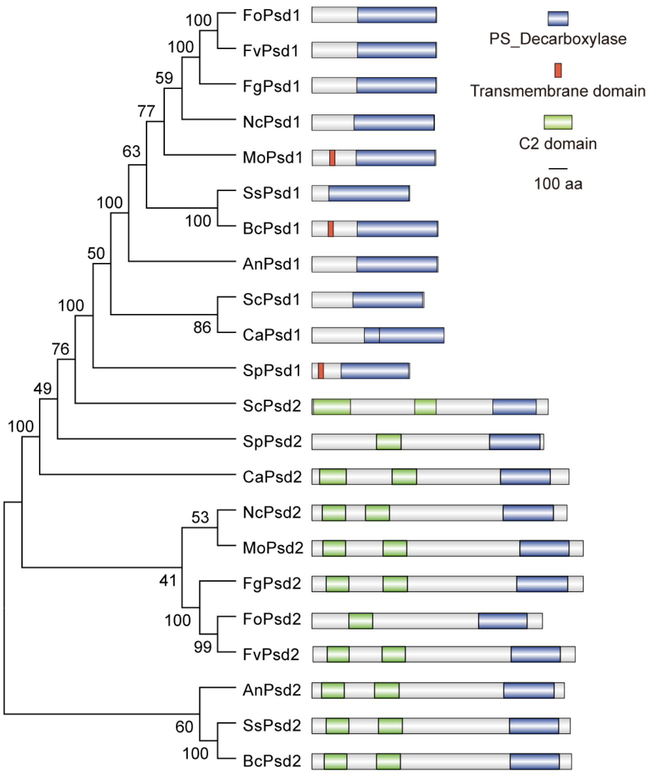


**Fig. S2** Phylogenetic and protein domain analyses of Psd homologs.

Values at the branching points of the phylogenetic tree represent the results of bootstrap analysis. An, *Aspergillus nidulans*; Bc, *Botrytis cinerea*; Ca, *Candida albicans*; Fg, *Fusarium graminearum*; Fo, *Fusarium oxysporium*; Fv, *Fusarium verticillioides*; Mo, *Magnaporthe oryzae*; Nc, *Neurospora crassa*; Sc, *Saccharomyces cerevisiae*; Sp, *Schizosaccharomyces pombe*; Ss, *Sclerotinia sclerotiorum*. Protein domains were predicted with SMART.

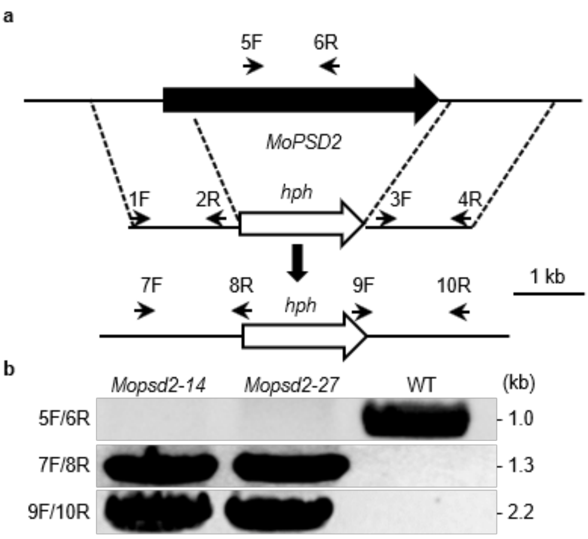


**Fig. S3** The *MoPSD2* gene replacement construct and mutants.

(**a**) The *MoPSD2* genomic region and the gene deletion construct. The upstream and downstream flanking fragments were amplified with primer pairs *PSD2*-1F/2R and *PSD2*-3F/4R, respectively, and ligated with the hygromycin B phosphotransferase gene (*hph*) cassette by double-joint PCR. Bar, 1 kb. (**b**) The *Mopsd2* mutants (*Mopsd2-14*, *Mopsd2-27*) were verified with primer pairs 5F/6R, 7F/8R and 9F/10R.

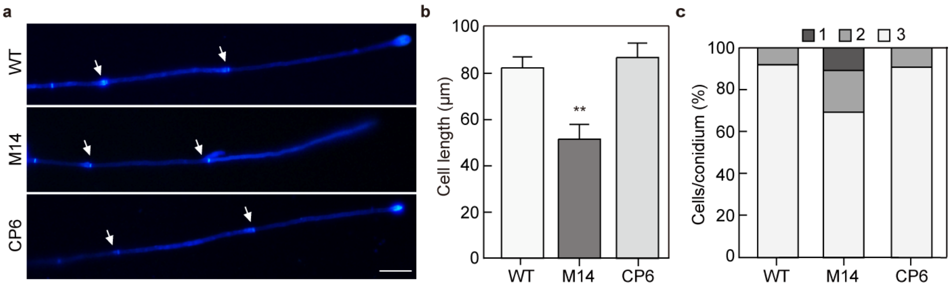


**Fig. S4** *MoPSD2* contributes to hyphal growth in *M. oryzae*.

(**a**) Hyphal tips of the wild-type (WT), *Mopsd2* mutant (M14) and *Mopsd2*/MoPsd2-GFP complementation (CP6) strains stained with Calcofluor white (CFW). Arrows indicate hyphal septa. Bar, 20 μm. (**b**) Cell length of the apical cells in the hyphal tips of strains. Error bars indicate standard deviations. Data were analyzed with Student's *t*-test. Asterisks represent significant differences (**$P < 0.05$). (**c**) Cell numbers in conidia of different strains.

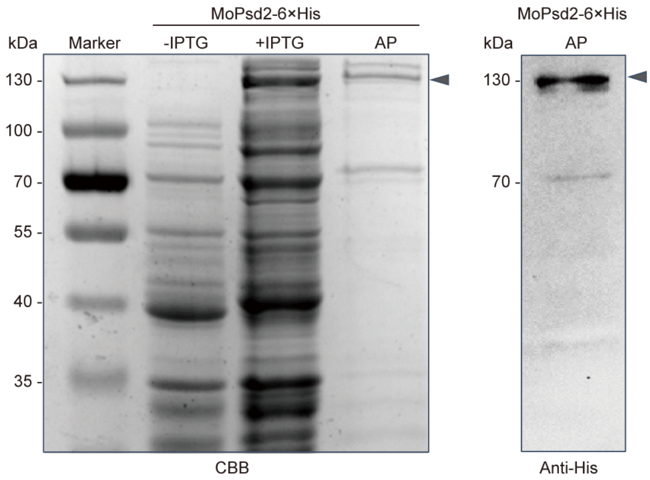


**Fig. S5** Purification and western blotting analysis of MoPsd2-6×His protein.
Coomassie brilliant blue (CBB) staining of total proteins isolated from *E. coli*. IPTG, isopropyl β-D-thiogalactoside; AP, affinity purification of proteins from the nickel column. Immunoblotting analysis of purified proteins with the anti-His antibody. Arrows indicate the MoPsd2-6×His protein.

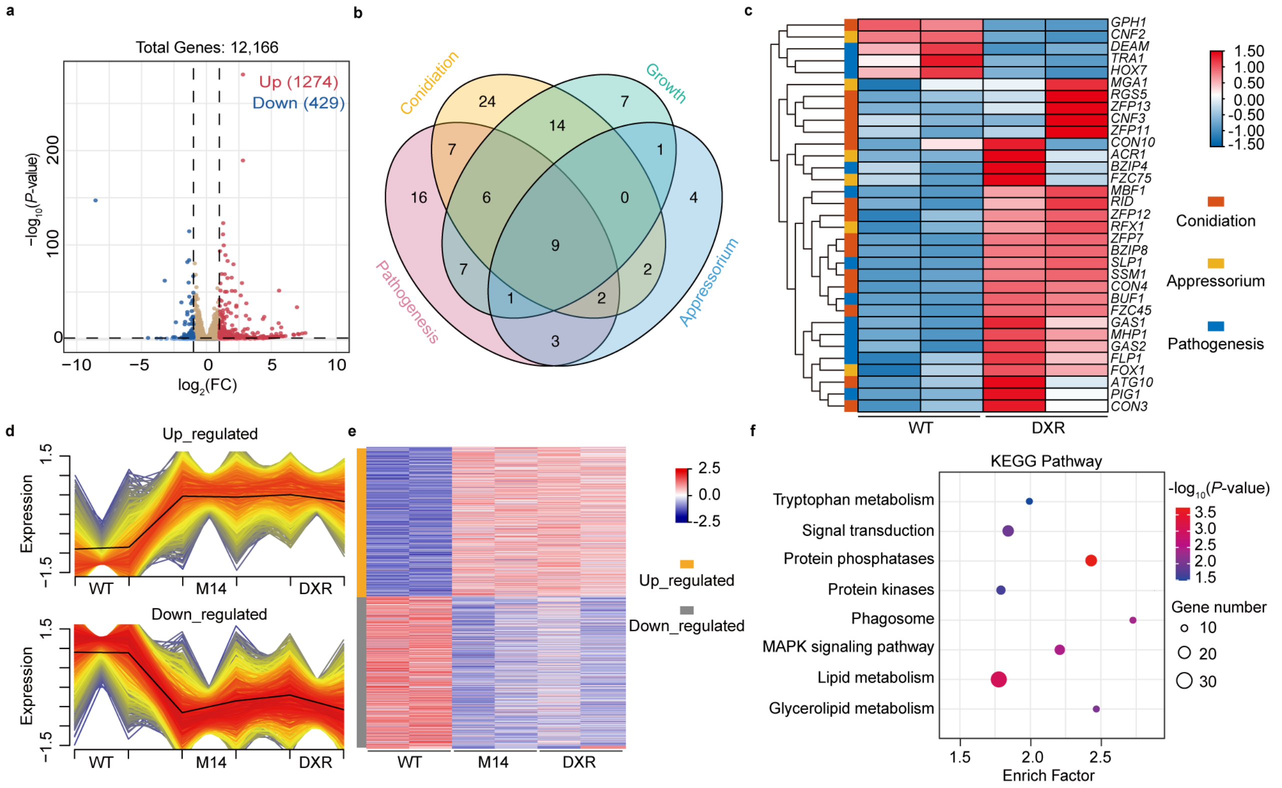


**Fig. S6** Doxorubicin treatment alters expression levels of genes involved in growth, conidiation, appressorium formation, and pathogenesis in *M. oryzae*.

(**a**) Volcano plot of differentially expressed genes (DEGs) in *M. oryzae* treated with or without doxorubicin. The x-axis shows the fold change (FC) of gene expression, while the y-axis indicates the significance. (**b**) Venn diagram of genes involved in development and plant infection. (**c**) Expression levels of genes involved in conidiation, appressorium formation and pathogenesis with or without doxorubicin. (**d**) Non-overlapping gene clusters derived from the *Mopsd2* mutant and *M. oryzae* cultured in media supplemented with or without doxorubicin. (**e**) Heat maps represent the degrees of similarity among the WT, *Mopsd2* mutant and doxorubicin treated samples. (**f**) KEGG functional enrichment analysis of genes from cluster 6 in (**e**).

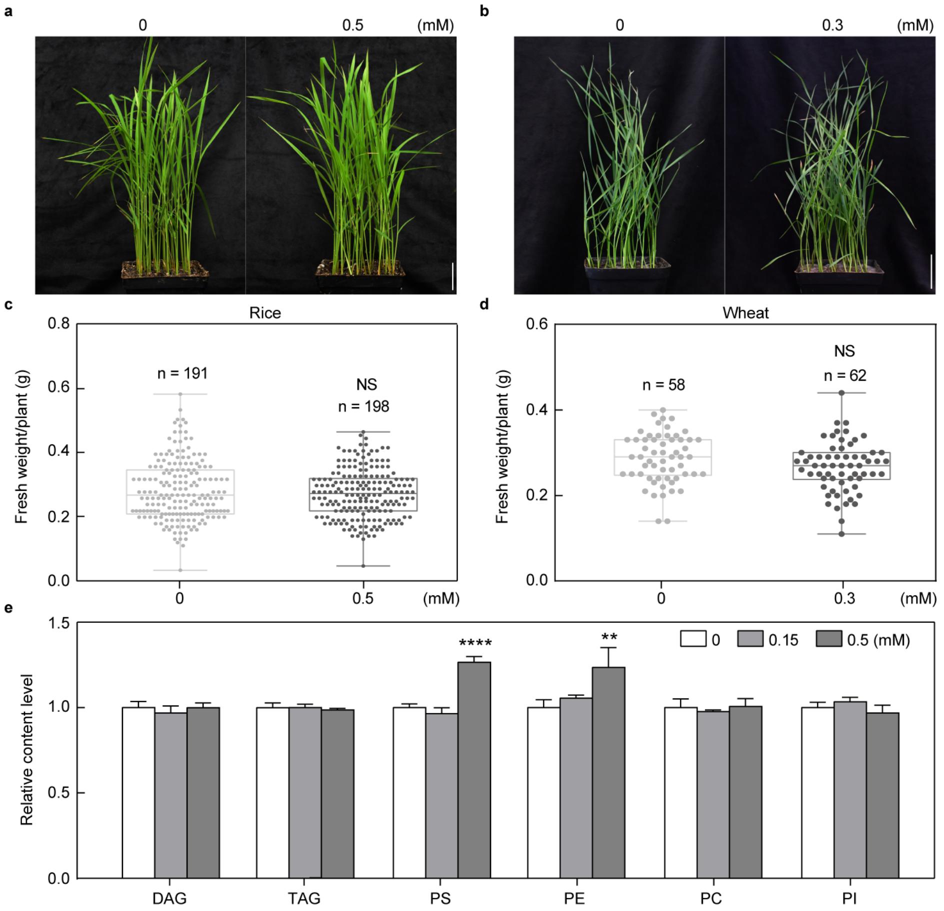


**Fig. S7** Phytotoxicity assays of doxorubicin on rice and wheat.

Rice (**a**) and wheat (**b**) seedlings were exposed to 0.15 mM doxorubicin for 7 and 5 days, respectively. Fresh weight per plant of rice (**c**) and wheat (**d**) was determined from three biological replicates. Data are displayed as box and whisker plots with individual data points: center line, median; box limits, 25th and 75th percentiles. (**e**) Lipidomics assays of rice samples treated with or without doxorubicin. Means and standard deviations were calculated from the number of individual plants as indicated. Error bars indicate standard deviations. Data were analyzed with Student's *t*-test. Asterisks represent significant differences (**$P < 0.01$, ****$P < 0.0001$); NS, not significant.

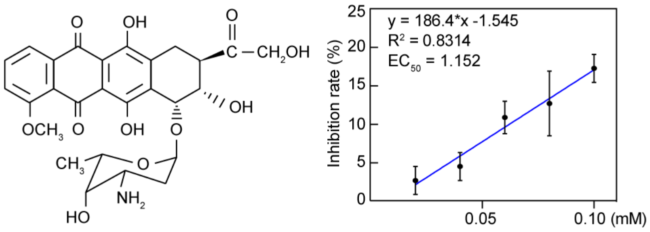


**Fig. S8** The concentration of doxorubicin for 50% of maximal effect (EC50).
Structure and EC50 values of doxorubicin. The EC50 of doxorubicin was calculated to be 1.152 mM based on the inhibition rate of mycelial growth on MM plates. Means and standard deviations (error bars) were calculated from three biological replicates.

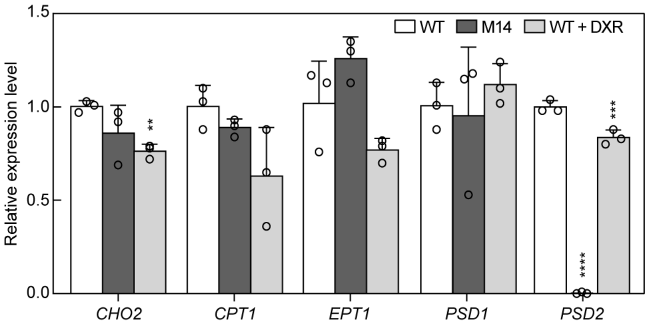


**Fig. S9** RT-qPCR assays of genes involved in PE biosynthesis in *M. oryzae*.

RT-qPCR assays of the genes in the *Mopsd2* mutant (M14) and the wild-type (WT) strain with or without doxorubicin (DXR). Means and standard deviations (error bars) were calculated from three biological replicates. Data were analyzed with Student's *t*-test. Asterisks represent significant differences (**$P < 0.05$, ***$P < 0.001$, ****$P < 0.0001$).

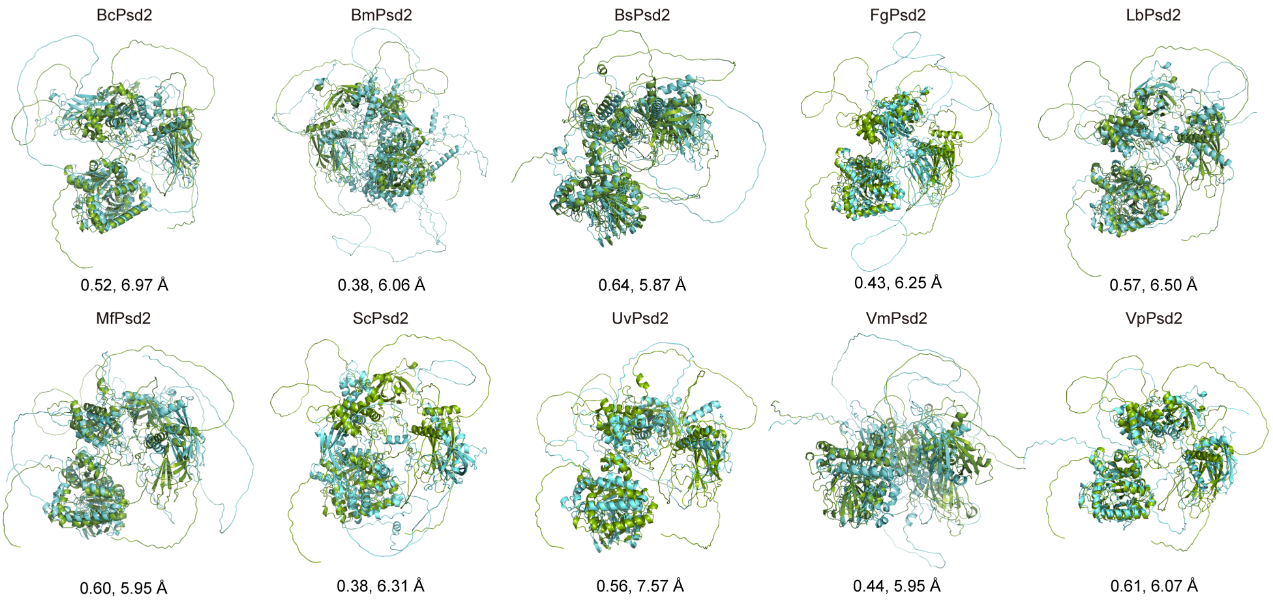


**Fig. S10** Predicted protein structures of Psd2 homologs of different fungal pathogens. Comparisons of Psd2 structures between *M. oryzae* and different fungal pathogens. The predicted structure of MoPsd2 is in green, and predicted structures of different Psd2 homologs in blue. Parameter 1, TM-score, template modeling score; parameter 2, RMSD, root mean square deviation. BcPsd2, XP024553066.1; BmPsd2, KAH7552113.1; BsPsd2, KAF5846341.1; FgPsd2, PCD31625.1; LbPsd2, KAH9877722.1; MfPsd2, KAG4031716.1; MoPsd2, XP003715218.1; ScPsd2, GAA23544.1; UvPsd2, GAO15339.1; VmPsd2, KUI72771.1; VpPsd2, KUI56681.1.
